# A bacterial lipid triggers membrane mechanosensing immunity in *Arabidopsis*

**DOI:** 10.64898/2026.04.23.720373

**Authors:** Marie-Dominique Jolivet, Michaela Neubergerová, Vaishali Gabani, Ana Álvarez-Mena, Axelle Grelard, Joris Sprakel, Birgit Habenstein, Marc Ongena, Magali Deleu, Roman Pleskot, Julien Gronnier

**Affiliations:** Plant Cell Biology, TUM School of Life Sciences, Technical University of Munich, Germany; Center for Plant Molecular Biology (ZMBP), University of Tübingen, Germany; Institute of Experimental Botany of the Czech Academy of Sciences, Prague, Czech Republic; Department of Experimental Plant Biology, Faculty of Science, Charles University in Prague, Prague, Czech Republic; University of Bordeaux, CNRS, Bordeaux INP, CBMN, UMR 5248, F-33600 Pessac, France; Laboratory of Biochemistry, Wageningen University and Research, Wageningen, The Netherlands; Microbial Processes and Interactions laboratory, TERRA Research Centre, Gembloux Agro-Bio Tech, University of Liège; Gembloux, 5030, Belgium; Laboratory of Molecular Biophysics at Interfaces, TERRA Research Centre, Gembloux Agro-Bio Tech, University of Liège; Gembloux, 5030, Belgium

## Abstract

The plant immune system engages cell-surface receptors that detect microbe-associated molecular patterns to initiate pattern-triggered immunity (PTI), and intracellular receptors that sense microbe-secreted effectors to activate effector-triggered immunity (ETI). Whether additional modes of microbial detection exist remains unclear. Here, we define membrane mechanosensing immunity (MSI), a third layer of immune signaling. A bacterial lipid, the main diffusible signal factor (DSF) from *Xanthomonas campestris pv. campestris*, acts as a membrane-active molecule that alters plasma membrane biophysical properties, activates *Mechanosensitive Channel of Small Conductance* (*MscS*)*-like* (*MSL*)-dependent immune signaling, and triggers a broad transcriptional reprogramming that overlaps with PTI and ETI. MSI modulates PTI signaling and requires both PTI and ETI components for effective disease resistance. These findings establish the sensing of metabolite-induced membrane perturbations as a mechanism of microbial detection.

## Main

Cellular organisms are continually exposed to potential microbial invasion, which has driven the evolution of sophisticated immune systems (*1–3*). In plants, the detection of microbes involves a two-tiered innate immune receptor system (*4*, *5*). At the cell surface, the perception of microbial-associated molecular patterns (MAMPs) by pattern recognition receptors (PRRs) leads to pattern-triggered immunity (PTI) (*6*). For instance, the *Arabidopsis thaliana* (hereafter Arabidopsis) leucine-rich repeat receptor kinase (LRR-RK) FLS2 perceives the flagellin epitope flg22 and forms a ligand-induced complex with its co-receptor BAK1 (also called SERK3) to initiate immune signaling events (*7*, *8*). Among them, the rapid MAPK phosphorylation, production of apoplastic reactive oxygen species (ROS) and calcium (Ca^2+^) influx are hallmarks of early PTI signaling (*9*, *10*). Intracellularly, the direct or indirect perception of microbial effectors by intracellular nucleotide-binding leucine-rich repeat (NLR) receptors induces effector-triggered immunity (ETI) (*11*, *12*). For instance, the bacterial effector protein AvrRps4 is recognized by a pair of Toll/interleukin-1 receptor/resistance protein (TIR)-NLRs, RRS1 and RPS4 (*13*). TIR-NLR signalling involves the lipase-like proteins EDS1, SAG101 and PAD4 (*14*) and helper NLRs of the ADR1 and NRG1 families (*15*). Both cell-surface and intracellular immune receptor systems modulate one another (*11*, *16–18*), exhibit direct molecular connections (*17*, *19*) and converge at membranes for signaling (*20*). Whether additional microbial detection systems contribute to plant immunity remains unclear. Diffusible signal factors (DSFs) are cis-2Δ unsaturated fatty acids acting as quorum sensing molecules to coordinate the behavior of bacterial communities (*21*). These molecules are produced by Gram-negative bacteria including several devastating phytopathogens such as *Xanthomonas campestris pv. campestris* (*Xcc*) and *Xylella fastidiosa*, as well as several human pathogens, including *Pseudomonas aeruginosa* and *Salmonella enterica* (*22*). In *Xcc*, the main DSF corresponds to a cis-2Δ-11-methyl-dodecenoic acid (hereafter referred to as DSF, Fig. 1A) which gradually accumulates and reaches micromolar ranges during infection (*23*). Its exogenous application has been reported to promote (*23*, *24*) or inhibit (*25*, *26*) immunity in Arabidopsis. Whether and potentially how DSF is perceived is unknown. Unlike flg22 perceived by FLS2, or the bacterial-derived medium-chain 3-hydroxy fatty acid (3-OH-FA) perceived by LORE (*27*, *28*), DSF treatment did not induce the rapid production of apoplastic ROS, MAPK phosphorylation, or Ca^2+^ influx in Col-0 (Fig. S1A-C). Similarly, DSF treatment did not induce ROS production in six additional ecotypes (fig. S1D), indicating that Arabidopsis does not encode a PRR that would enable the detection of DSF and the activation of PTI.

**Figure 1.**
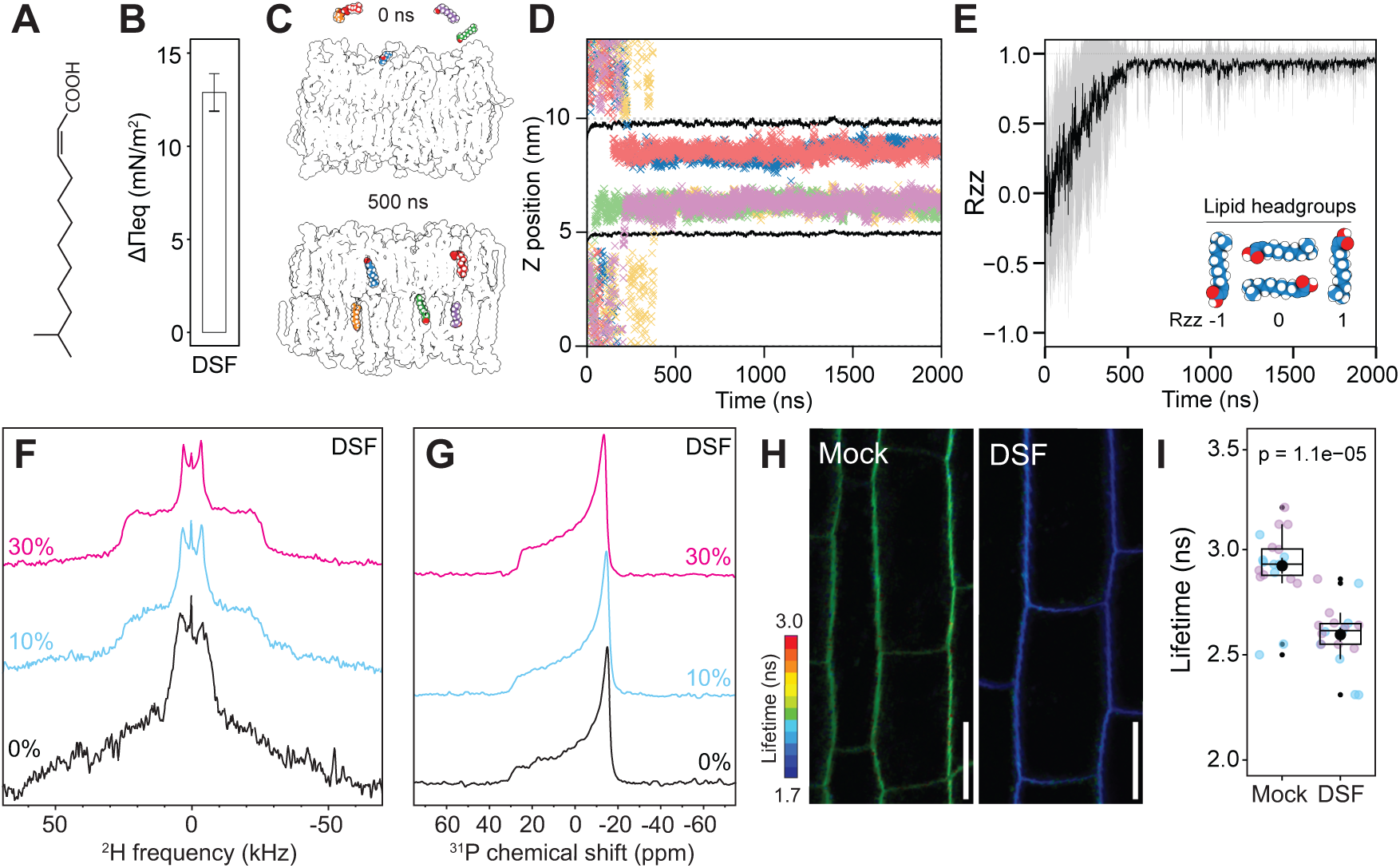
DSF perturbs plasma membrane biophysics. (**A**) Chemical structure of DSF. (**B**) Mean ± SE of surface pressure variation at the equilibrium (ΔΠeq) upon adsorption of DSF to a PLPC/sitosterol/glucosylceramide (GluCer) (6:2:2 molar ratio) monolayer from 3 independent experiments. (**C**-**E**) Molecular dynamic simulations of DSF insertion in model plant PM outer leaflet. Side view images of representative time points 0 and 500 ns. (C), Z position of DSF center of mass over the time for one selected simulation repeat. Each cross represents an individual DSF molecule at each time point. Different colors represent five different DSF molecules, colors correspond to (C). Average z position of the phosphorus headgroup atoms is shown as a black line. Remaining 2 repeats are shown in Fig. S2. (D) Analysis of the rotation of the DSF molecule relative to the Z-axis (Rzz). The DSF carboxyl group predominantly faces the lipid headgroups, while the acyl chain aligns with the acyl chains of phospholipids and sphingolipids. Mean Rzz of 15 DSF molecules (from three simulation repeats) shown as solid line, standard deviation shown in gray. (E). (**F-G**) Solid-state NMR spectroscopy of POPC-d31/sitosterol/GluCer (7:1:2 molar ratio) liposomes in the absence (black) or in presence of 10 % (light blue) or 30 % (pink) DSF at 263 K, (F) ^2^H quadrupolar spin echo spectra, (G) ^31^P Hahn echo spectra. (**H-I**) Representative FLIM images (H) and lifetime analysis (I) of di-4-ANEPPDHQ in root epidermal cells treated with 10 µM DSF or mock (DMSO) for 30 min. Each data point represents lifetime (τ1) per cell from at least 9 seedlings; colors indicate independent experiments. *P* value reports a Wilcoxon test. Scale bar = 20 µm.

### DSF inserts in model membranes and perturbs membrane biophysics

Because DSF is a lipid, we hypothesized that it inserts into membranes. In agreement, in Langmuir trough assays we observed that DSF increased surface pressure (Fig. 1B and fig. S2A), indicative of its incorporation into a model lipid monolayer *in vitro*. To obtain a molecular view of DSF insertion into a model plant plasma membrane outer leaflet (composition detailed in Table 1) we performed all-atom molecular dynamics simulations. DSF consistently inserted into the bilayer within the first 500 ns (Fig. 1, C and D, and fig. S2B). In the membrane, the carboxyl group of DSF oriented toward lipid headgroups (Fig. 1E), while its acyl chain interacted with phospholipid and sphingolipid acyl chains, with a moderate preference for unsaturated chains (fig. S2D) but no specificity for lipid species (fig. S2C). DSF exerted two opposing effects: it increased acyl-chain order parameters, suggesting a local increase in membrane rigidity, while enhancing the motility of phospholipid headgroups (fig. S2, E to I). We next tested whether DSF alters membrane properties *in vitro* using solid-state nuclear magnetic resonance (ssNMR) spectroscopy on liposomes containing deuterated phosphatidylcholine (POPC-d31) as a probe. Near the fluid-to-gel phase transition temperature, ^2^H NMR spectra revealed a dose-dependent stabilization of the liquid-disordered phase by DSF, reflected in narrower quadrupolar splittings and a reduced first spectral moment M_1_ (Fig. 1F and fig. S3A) (*29*). This effect extended to lipid headgroups, as the ^31^P spectra showed a reduced chemical shift anisotropy linewidth (Fig. 1G and fig. S3B). The local acyl-chain ordering observed in simulations together with the enhanced ensemble fluidity measured by ssNMR indicates that DSF induces spatially heterogeneous membrane states, in which lipid perturbations generate non-uniform packing. Altogether, these results show that DSF inserts into model membranes and affects membrane biophysical properties. To assess whether DSF similarly affects plasma membrane properties *in vivo*, we used the solvatochromic dye di-4-ANEPPDHQ to measure membrane order (*30*, *31*). Both ratiometric and lifetime imaging revealed that DSF treatment increased plasma membrane disorder (Fig. 1, H to I, and fig. S4, A and B). Consistently, the molecular rotor N⁺-BODIPY (*32*), showed a reduced fluorescence lifetime upon DSF treatment, further indicating increased plasma membrane disorder (fig. S4, C to D). These observations demonstrate that DSF perturbs plasma membrane properties *in vivo*.

**Table 1.** Lipid composition of the model membranes used in molecular dynamics simulations.

| Lipid | Saturation | % |
| --- | --- | --- |
| **Simulations of Fig. 1 C-E and fig. S2 B** |  |  |
| POPC | 16:0/18:1 | 14 |
| PLPC | 16:0/18:2 | 14 |
| POPE | 16:0/18:1 | 5 |
| PLPE | 16:0/18:2 | 5 |
| OEPS | 18:1/22:1 | 3 |
| PLPG | 16:0/18:2 | 1 |
| POPI | 16:0/18:1 | 1 |
| Sitosterol |  | 29 |
| GIPC (Man(a1-4)-GlcA(a1-2)-IPC) | d18:1/h24:0 | 20 |
| Glucosylceramid | d18:1/h24:1 | 5 |
| Ceramid | d18:1/h24:0 | 3 |
| **Simulations of fig. S2 C-I** |  |  |
| POPC | 16:0/18:1 | 70 |
| Sitosterol |  | 10 |
| GIPC (Man(a1-4)-GlcA(a1-2)-IPC) | d18:1/h24:0 | 20 |

### DSF induces *MSL-*dependent signaling

We hypothesized that the changes in plasma membrane order induced by DSF may activate mechanosensitive channels. Among the mechanosensitive channels encoded in Arabidopsis, three families present plasma membrane-localized members (*33*), the Ca^2+^-permeable Mid1-Complementing Activity (MCA) 1 and 2 (*34*, *35*), the Ca^2+^-permeable OSCAs/TMEM63 (*10*, *36*, *37*) and the anion-permeable Mechanosensitive channel of small conductance (MscS)-like (MSL) (*38–40*). In agreement with the Aequorin-based assays (fig. S1B), no changes in intracellular Ca^2+^ concentrations were captured using the R-GECO1 sensor (fig. S1E), suggesting that DSF-induced changes in membrane properties do not gate MCAs or OSCAs channels. Single-channel patch clamp electrophysiology analyses of MSL10 in oocytes indicate that MSL preferentially mediates chloride (Cl^-^) efflux (*39*). Using the E^2^GFP reporter (*41*, *42*), we observed that DSF treatment induced a decrease in intracellular Cl^-^ (Fig. 2, A and B), suggestive of Cl^-^ efflux. Furthermore, the loss of the five-plasma membrane localized MSLs, (*msl4/5/6/9/10* (*38*), hereafter referred to as *msl-q*) abolished the decrease in intracellular Cl^-^ induced by DSF (Fig. 2A and B), suggesting that DSF gates MSLs. We next asked whether DSF treatment was associated with additional *MSL*-dependent intracellular signaling events. Consistent with 2′,7′-dichlorofluorescein diacetate staining (*24*), the genetically-encoded reporter HyPer7 revealed that DSF induced the accumulation of intracellular ROS (iROS) in wild type, whereas this response was abolished in *msl-q* (Fig. 2, C and D). Late cellular responses were also investigated. Using the voltage-sensitive dye DIBAC(4)_3_, we observed that 24-hour DSF treatment induces a depolarization of the plasma membrane in an *MSL*-dependent manner (Fig. 2, E and F). Within this time frame, an increase in cytosolic Ca^2+^ levels was also measured by R-GECO1 (Fig. 2, G and H) and by the Aequorin reporter (fig. S5, A and B). Finally, 24-hour DSF treatment led to an increase in MAPK phosphorylation (Fig. 2I) and to a sustained iROS accumulation, both of which were abolished in *msl-q* (fig. S5, C and D). Altogether, these observations show that DSF induces early and late *MSL*-dependent intracellular signaling events.

**Figure 2.**
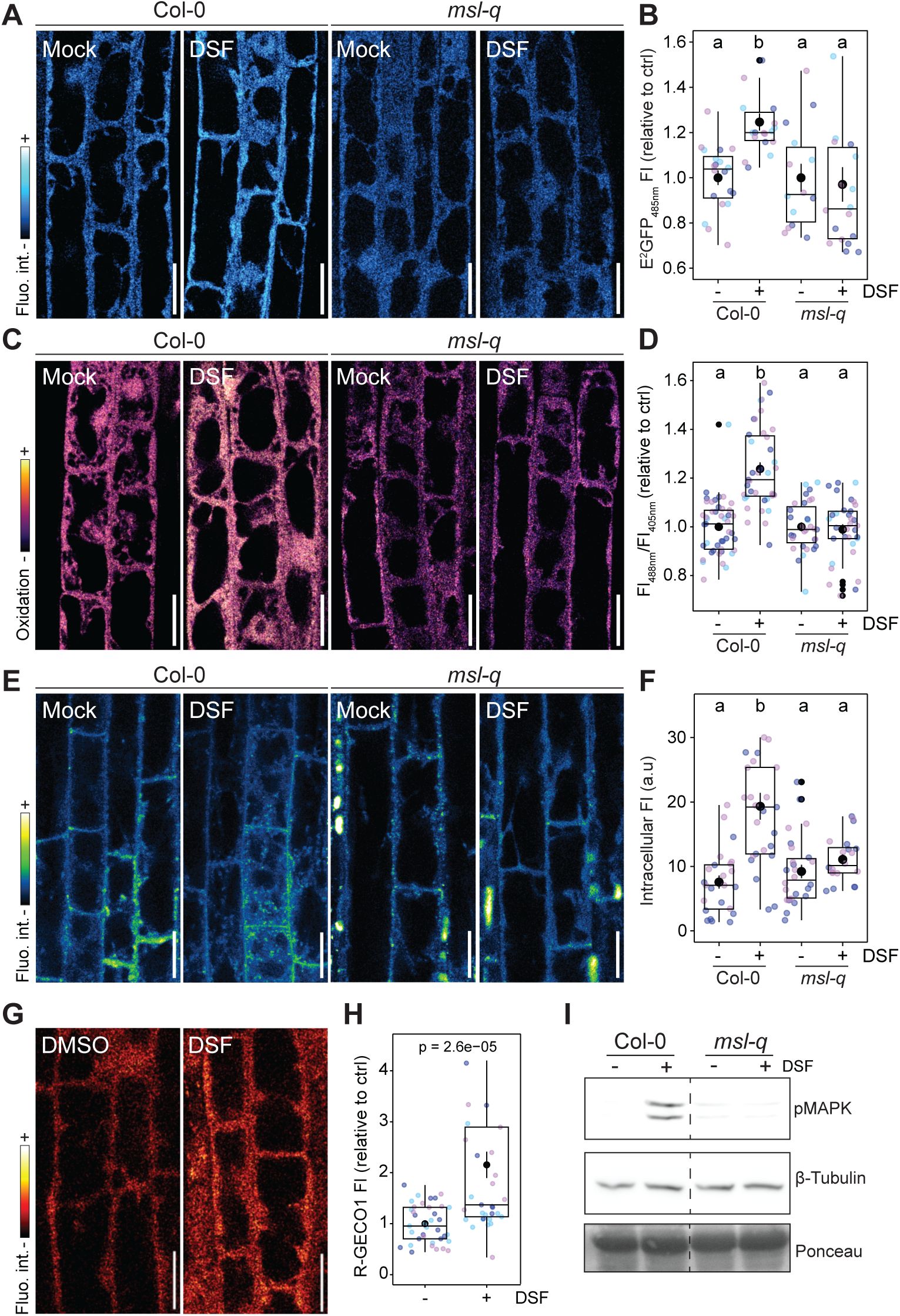
DSF induces mechanosensitive channel-dependent cell signaling. (**A**-**B**) Analysis of intracellular Cl^-^. Representative confocal images (A) and analysis of fluorescence intensity (B) of E_2_GFP in Col-0/R-GECO1-GSL-E^2^GFP and *msl-q*/R-GECO1-GSL-E^2^GFP root epidermal cells treated with 10 µM DSF or mock (DMSO) for 30 min. Each data point represents mean fluorescence intensity of an individual cell; colors indicate independent experiments with at least 14 seedlings analyzed per condition. (**C**-**D**) Analysis of intracellular ROS. Representative confocal images (C) and ratiometric analysis of fluorescence intensity (D) of Hyper7 in root epidermal cells treated with 10 µM DSF or mock (DMSO) for 30 min. Each data point represents mean fluorescence of an individual cell; colors indicate independent experiments with at least 17 seedlings analyzed per condition. (**E**-**F**) Analysis of plasma membrane depolarization. Representative confocal images (E) and analysis of intracellular fluorescence (F) of DiBAC_4_(3) in root epidermal cells treated with 10 µM DSF or mock (DMSO) for 24 h. Each data point represents mean intracellular fluorescence of a cell; color indicates an independent experiment, with at least 11 seedlings analyzed per condition. (**G**-**H**) Analysis of intracellular Ca^2+^. Representative confocal images (G) and analysis of fluorescence intensity (H) of R-GECO1 in Col-0/R-GECO1-GSL-E^2^GFP in root epidermal cells treated with 10 µM DSF or mock (DMSO) for 24 h. Each data point represents mean fluorescence of an individual cell; color indicates an independent experiment with at least 10 seedlings analyzed per condition. *P* value reports a Student’s t-test. (**I**) Western blot analysis of MAPK phosphorylation 24 h after 10 µM DSF treatment in 10-day-old seedlings. α-p44/42-ERK antibodies were used to probe phosphorylated MAPK. β-Tubulin immunoblotting and Ponceau staining were used to assess loading. In (B), (D) and (F), letters indicate statistically significant differences between conditions (one-way ANOVA followed by a Tukey’s HSD; p<0.01). In (A), (C), (E) and (G), scale bar = 20 µm.

### DSF induces *MSL-*dependent immunity that requires both ETI and PTI

We next investigated whether DSF-induced signaling is linked to immunity. RNA-sequencing revealed that DSF induced a vast transcriptional reprogramming, with 4361 differentially expressed genes (DEGs) identified throughout the experiments (log2(fold change, FC)>1, Padj<0.05) (fig. S6, A and B). We compared these transcriptional responses with MAMP-induced PTI responses (*43*) and the ETI transcriptional response induced by the bacterial effector protein AvrRps4 (*16*). We found that DSF-induced DEGs overlap with both MAMP-induced DEGs (described as PTI response) and with AvrRps4-induced DEGs (described as ETI response) (Fig. 3, A and B). Furthermore, a large set of DSF-induced DEGs overlaps with all MAMP-induced DEGs (fig. S6C), indicating that DSF induces an immune transcriptional response. Consistently, DEGs commonly induced by DSF, PTI and ETI are predominantly enriched for immune-related Gene Ontology terms (fig. S7). Interestingly, DSF regulated common and distinct gene family members commonly associated with immune signaling (fig. S6D) and hormonal responses (fig. S6E). We conclude that DSF induces a broad immune transcriptional response that overlaps with PTI and ETI transcriptional responses. We next asked whether DSF-induced cellular responses were associated with an effective immune response. To this end, we tested infection with the bacterium *Pseudomonas syringae* pv. tomato DC3000 (Pto), which, unlike *Xcc,* does not utilize DSF as a quorum-sensing molecule. As shown for *Xcc* (*24*), DSF pre-treatment led to an increase in disease resistance against Pto (Fig. 3C). In contrast, *msl-q* was unable to mount an effective DSF-induced resistance and presented an increased sensitivity to Pto infection in control condition (Fig. 3C). Consistent with previous reports linking MSLs with immunity (*44*, *45*), we conclude that *MSLs* are components of the plant immune system and that DSF-induced and *MSL*-dependent signaling events culminate in effective immune responses promoting disease resistance against Pto. We next asked about potential functional connections between DSF-induced immunity and PTI or ETI. We observed that flg22-induced MAPK phosphorylation and flg22-induced ROS production were not affected in *msl-q* (fig. S8, A and B). In addition, *MSLs* were dispensable for flg22-induced disease resistance (fig. S8C), indicating that MSLs do not correspond to core PTI components. Furthermore, DSF did not trigger cell death at the concentration tested (5-40 µM), a cellular response commonly associated with ETI. Interestingly, 24 h DSF pre-treatment promoted flg22-induced ROS production in an *MSL-*dependent manner (Fig. 3D) indicating that DSF-induced signaling promotes PTI. In addition, as shown for AvrRps4-induced ETI (*16*), the immune loss-of-function *BAK1* allele *bak1-5* (*46*) abolished DSF-induced resistance (Fig. 3E). Further, DSF-induced resistance was impaired in the NLR *helperless* mutant (*15*) (Fig. 3E), indicating that DSF-induced resistance relies on both ETI and PTI signaling modules.

**Figure 3.**
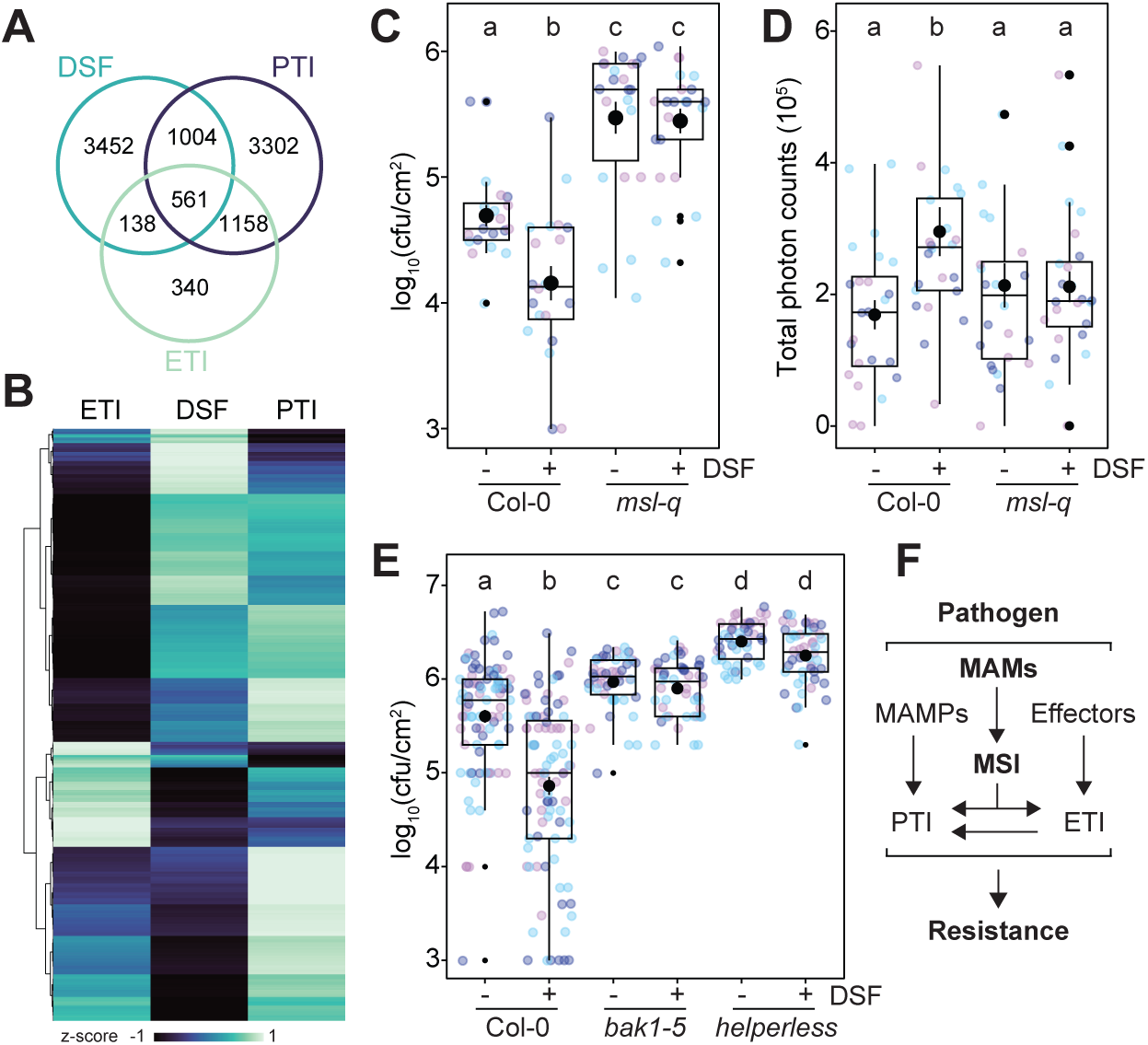
DSF induces *MSL-*dependent immunity that requires both ETI and PTI. (**A**) Venn diagram analysis of shared and treatment specific DEGs (|log2(FC)| > 1; Padj< 0.05) induced by DSF, MAMPs (PTI), and AvrRps4 (ETI). (**B**) Heatmap of DEGs induced by DSF, MAMPs (PTI), and AvrRps4 (ETI), normalized as z-scores. The genes are hierarchically clustered according to their behaviour across conditions. (**C**) Number of *Pto* DC3000 bacteria 3 days after inoculation of plants pre-treated with 5 µM DSF or mock (DMSO) for 24 h. Each data point represents an individual leaf; their color indicates an independent experiment. (**D**) Apoplastic ROS production after 100 nM flg22 in plants pretreated overnight with 5 µM DSF or mock (DMSO). Each data point represents an individual leaf disc; colors indicate independent experiments with 24 leaf discs analyzed per condition. (**E**) Number of *Pto* DC3000 bacteria 3 days after inoculation of plants pre-treated with 5 µM DSF or mock (DMSO) for 24 h. Each data point represents an individual leaf; colors indicate independent experiments with at least 12 plants analyzed per condition. (**F**) Proposed model of the contribution of MSI in plant immunity. In (C), (D) and (E), letters indicate statistically significant differences between conditions (one-way ANOVA followed by a Tukey’s HSD; p<0.05).

### Membrane-active molecules activate mechanosensitive immunity

The plant immune system is commonly defined as a two-tiered innate immune receptor system. Our findings define a mode of microbial detection that departs from receptor-based perception and relies instead on the sensing of membrane perturbation. We show that DSF behaves as a membrane-active molecule (MAM) that inserts into the plasma membrane, alters its physical properties, and activates mechanosensitive MSL channels, thereby triggering membrane mechanosensing immunity (MSI) (Fig. 3F). Our study further shows that MSI operates in parallel with PTI and ETI and converges with these pathways to mount an effective immune response. ETI has been shown to potentiate PTI components, and both ETI and PTI coordinate the induction of strong immunity against bacterial pathogens (*16*, *18*). Here we show that DSF-induced MSI requires core components of both PTI and ETI (Fig. 3E), and that DSF-induced MSI primes flg22 responsiveness (Fig. 3D). Together, these findings support a multilayered immune system in which MSI, PTI, and ETI collectively shape robust defense responses (Fig. 3F).

High concentrations of DSF (40 µM) have also been reported to inhibit immunity (*26*). Consistently, we observed that elevated concentrations of DSF (40 µM) inhibited flg22-induced ROS and promoted disease (fig. S9, A-C) and that these effects occurred in an *MSL*-dependent manner (fig. S9, A to C). DSF did not however affect flg22-induced FLS2-BAK1 complex formation, MAPK phosphorylation (fig. S9D) or Ca^2+^ influx (fig. S9, E and F), suggesting that hyperactivation of *MSL*-dependent signaling by DSF specifically dampens the activation of NADPH oxidases by flg22. Interestingly, the NADPH oxidase RBOHD is required for microbiota homeostasis, and loss of *RBOHD* particularly favors the proliferation of Xanthomonadales (*47*, *48*). During infection, DSF gradually accumulates and can reach substantial levels (40-100 µM) (*23*), Thus, *Xcc* may have co-opted the inhibitory effect of high DSF concentration on PTI signaling to promote its proliferation. These findings suggest that MSI behaves as a tunable module whose output depends on the amplitude or regime of membrane perturbation. Such a model reconciles previously conflicting reports on DSF and implies that pathogens may exploit high concentrations of MAMs to reprogram host signaling through the same pathway that, at lower levels, contributes to defense. In animals, the mechanosensitive channel PIEZO1 participates in immunity and senses pathogen-induced membrane deformation (*49–51*). Our study expands the role of mechanosensitive channels to the detection of microbial metabolites in Arabidopsis and supports a unifying framework in which mechanosensitive channels translate membrane perturbations into immune responses. In this model, membranes function as biophysically encoded interfaces for immune perception, where mechanosensitive channels decode perturbations associated with microbial colonization to initiate immune signaling. This framework may also at least partly underlie for the gating of plant defenses in response to pathogen-derived (*52*), damage- (*53*) and raindrop-induced (*54*) mechanical cues. Several important bacterial pathogens of plants and animals produce DSF-family molecules that have been linked to immunity (*22*) and may similarly activate mechanosensitive channels. We recently showed that membrane remodeling induced by the *Bacillus* lipopeptide surfactin (Srf) triggered mechanosensitive channels-dependent signaling and disease resistance (*55*). The ability of the structurally divergent MAMs such as DSF and Srf to trigger MSI, together with the emerging conserved role of mechanosensitive channels in immunity, suggests that a wide range of microbial-derived compounds may function as MAMs and induce immune responses across kingdoms.

## Acknowledgements

We thank all members of the NanoSignaling Laboratory for fruitful discussions and comments on the manuscript. We thank Rainer Waadt for providing the R-GECO1-GSL-E^2^GFP plasmid and the Col-0/R-GECO1-GSL-E^2^GFP transgenic line, Alexandre Martiniere for providing the Hyper7 plasmid and the Col-0/Hyper7 transgenic line, Cyril Zipfel for proving the *bak1-5* mutant, Farid El Kasmi for providing the *helperless* mutant and Sebastian Pfeifmeir for providing the Pto DC3000 bacterial strain. Bio-imaging was performed at the ZMBP microscopy facility of the University of Tübingen and at the Center for Advanced Light Microscopy (CALM) of the TUM School of Life Sciences.

## Funding

This research was funded by the University of Tübingen (to J.G.), the Technical University of Munich (to J.G.), the Deutsche Forschungsgemeinschaft (DFG) grant B01-TRR356 (to J.G.), the Belgian Funds for Scientific Research (PDR Project T.0081.24) to M.D., the Feder Programme 2021-2027, the project PHENIX Biocontrol ULiege-SPW to M.D. and M.O, and the French National Research Agency ANR (BH, grant number ANR-23-CE11-0005-01). Computational resources used for molecular dynamics simulations were provided by the e-INFRA CZ project (ID: 90254), supported by the Ministry of Education, Youth and Sports of the Czech Republic.

## Author contributions

Conceptualization M-D.J. and J.G.; project administration, J.G; investigation, M-D.J., M.N., A.A-M., A.G.; funding acquisition, J.G.; resources, J.S., V.G. and M.O.; supervision, J.G., R.P., B.H., and M.D.; writing – original draft, M-D.J. and J.G.; writing – review and editing, M-D.J. M.N., A.A-M., A.G., J.S., B.H., M.O., M.D., R.P. and J.G.

## Competing interests

The authors declare no competing interests.

## Data and materials availability

All data that support the findings of our study are available in the manuscript or supplementary materials.

## Material and Methods

### Plant material and growth conditions

The *Arabidopsis thaliana* ecotype Columbia-0 (Col-0) was used as the wild-type control. For experiments on seedlings, Arabidopsis seeds were surface sterilized using 70% ethanol, stratified for 24 hours at 4 °C, sown onto solid ½-strength Murashige and Skoog (MS) medium (1 % sucrose, 0.8 % agar, pH 5.7) and grown at 20-22 °C with a 16-hour photoperiod for 5 days. For experiments using adult plants, Arabidopsis seeds were surface sterilized using 70% ethanol, stratified for 24 hours at 4 °C, sown in soil and grown at 20-22 °C with a 10-hour photoperiod for 4 to 5 weeks. The Arabidopsis *msl4/5/6/9/10* (*38*), *moca1* (*56*), *bak1-5* (*46*), *helperless* (*15*), Col-0/HyPer7 (*57*), and Col-0/R-GECO1-GSL-E^2^GFP (*42*) were previously described. The *msl4/5/6/9/10* (*msl-q)* mutant was transformed with R-GECO1-GSL-E^2^GFP (*42*) to obtain *msl-q/*R-GECO1-GSL-E^2^GFP and with HyPer7 (*57*) to obtain *msl-q*/HyPer7.

### Chemicals and peptides

DSF (Merck, 42052) was dissolved in DMSO at a stock concentration of 100 mM. Lovastatin (Merck, PHR1285) was dissolved in DMSO at a stock concentration of 1 mM. Flg22 (QRLSTGSRINSAKDDAAGLQIA, Synpeptide) was dissolved in sterile double-distilled water at a stock concentration of 10 mM. 3-OH-FA (Merck, H3648) was dissolved in DMSO at a stock concentration of 100 mM. For treatment, the stock solutions were diluted in the appropriate medium and the concentrations specified in each figure.

### Molecular dynamics

All molecular dynamics (MD) simulations were performed using GROMACS (*58*) with the CHARMM36m force field (*59*). DSF was parameterized using ParamChem, accessible via the CHARMM-GUI web server (*60*).The SDF files for parametrization were obtained from PubChem (Conformer3D_COMPOUND_CID_11469920). For the simulations with model plant plasma membrane outer leaflet a 10.6 × 10.6 nm lipid bilayer was constructed using the CHARMM-GUI web server (*60*, *61*). The lipid composition used was selected according to (*62*) and is detailed in Table 1. Five DSF molecules were added in close proximity to the membrane in a 10.6 × 10.6 × 12.5 nm simulation box. The system was solvated with TIP3P water and 0.05 M CaCl₂, and neutralized. The multi-site Ca^2+^ (CAM) parameters were applied (*63*). The system was energy minimized using the steepest descent algorithm (up to 5000 steps), followed by 3750 ps equilibration with position restraints on DSF and lipids. Three 2000 ns production runs starting from different initial velocities were performed using a 2-fs time step in the NPT ensemble. Pressure was maintained at 1 bar using the Parrinello-Rahman barostat (coupling constant: 5.0 ps, compressibility: 4.5 × 10⁻⁴). Temperature was set to 303.15 K and regulated using the V-rescale thermostat (coupling constant: 1.0 ps). Z positions were analysed using MDAnalysis (*64*). Rotation relative to z axis (Rzz) and radial distribution function (RDF) were calculated using Gromacs built-in tools *gmx rotmat* and *gmx rdf.* RDF was calculated over the last 250 ns of the simulation. ChimeraX was used for the visualizations (*65*, *66*).

For the simulations with POPC/sitosterol/glycosylinositol phosphorylceramide (GIPC) a 8 x 8 nm membrane was built using the CHARMM-GUI web server (*60*, *61*) and positioned in a 8 x 8 x 12 nm simulation box. The system was solvated with TIP3P water and 0.05 M CaCl₂, and neutralized. The multi-site Ca^2+^ (CAM) parameters were applied (*63*). Similarly, membranes in which a portion of POPC, sitosterol and GIPC was replaced by 10% or 30% DSF were built. Energy minimization, equilibration and three 500 ns production runs starting from different initial velocities were performed using the same simulation parameters as described above. Acyl chain order was analysed using gorder tool (*67*). Motion of phospholipid headgroups was analysed using Gromacs built-in tool *gmx gangle.* For POPC, the angle was defined between the vector connecting the phosphorus and nitrogen atoms and the membrane normal (z-axis).

### Langmuir trough-based adsorption experiments

Adsorption experiments at constant surface area were performed in a KSV (Helsinki, Finland) Minitrough (190 cm^3^) equipped with a Wilhelmy plate. The lipid solution (PLPC:sitosterol:GluCer, 6:2:2 molar ratio) was spread at the air-subphase (10 mM MES, 150 mM NaCl, pH 5.8) interface to reach an initial surface pressure of 15.0 ± 0.4 mN/m^2^. After a 15-min wait for solvent evaporation and film stabilization, 20 µL of DSF (48 mM, dissolved in DMSO) was injected into the subphase, underneath the pre-formed lipid monolayer to a final concentration of 12 μM. The adsorption of DSF to the lipid monolayer was monitored by the increase of surface pressure. The subphase was continuously stirred, and temperature was maintained at 20 ± 1°C. No change in surface pressure was observed after injection of DMSO.

### Preparation of DSF-containing lipid vesicles for solid-state NMR

POPC, POPC-d31, sitosterol and GluCer were obtained from Avanti Polar Lipids, Inc. (USA). For sample preparation, the DSF compound (Merck, 42052) was dissolved together with lipids (POPC-d31:sitosterol:GluCer, molar ratio 7:1:2) in a chloroform/methanol mixture (2:1, v/v), maintaining a DSF-to-lipid molar ratio of 10 or 30%. The organic solvents were evaporated under a gentle stream of air. The resulting lipid film was then hydrated and subsequently lyophilized overnight. The resulting powder was resuspended in 100 µL of deuterium-depleted water and homogenized through three free-thaw cycles involving vortexing, rapid freezing in liquid nitrogen (−196 °C, 1 min) and incubation in a 40 °C water bath for 10 min. The same protocol was used for the control sample, prepared under identical buffer conditions in the absence of DSF. The resulting liposome suspensions were turbid, indicating particle sizes in the range of approximately 0.1 – 1 µm.

### 31P#and ^2^H wide-line solid-state NMR spectroscopy

^2^H static solid-state NMR experiments were performed using a phase-cycled quadrupolar spin-echo pulse sequence at a resonance frequency of 76.8 MHz on a Bruker Avance III 500 MHz spectrometer equipped with a 4 mm double-resonance HX CP-MAS probe. A 90° pulse length of 3.25 µs was used, with an interpulse delay of 50 µs and a recycle delay of 2 s. Spectra were acquired over a spectral width of 500 kHz with a number of scans between 1k to 4K. Measurements were carried out at temperatures ranging from 263 to 313 K, with samples equilibrated for 20 min at the target temperature prior to acquisition. Data processing was performed using TopSpin 4.0.6 (Bruker), applying a line broadening of 500 Hz before Fourier transform. First-order spectral moments (M1) were calculated using NMRDepaker (provided by Dr. Sébastien Buchoux), and simulations were carried out with NMR-099 (provided by Arnaud Grélard) to determine local order parameters (|2S_CD_|) along the acyl chains of POPC-d31(*29*, *68*). ^31^P NMR experiments were carried out on a Bruker Avance III HD spectrometer operating at the proton frequency of 400 MHz. Spectra were acquired under static conditions using a Hahn spin-echo sequence with proton decoupling during the acquisition. 1200 to 2k scans were collected depending on the sample. The 90° pulse length was 8 µs, with an echo delay of 40 µs and a recycle delay of 5 s. The spectral width was set to 64 kHz. All data were processed using TopSpin 4.0.6 (Bruker), applying 200-line broadening to the free induction decay prior to Fourier transformation (*69*, *70*).

### DIBAC_4_(3) staining and imaging

DIBAC_4_(3) staining and imaging were performed as previously described (*55*). Five-day-old seedlings were stained for 15 min with 10 µM DiBAC_4_(3) (Thermo Fisher Scientific, B438), washed twice with liquid ½ MS medium and mounted in liquid ½ MS medium between a microscope slide and coverslip for imaging. Roots were imaged on a Leica SP8 confocal microscope equipped with a 40x water-immersion objective (NA 1.2). DiBAC_4_(3) was excited at 514 nm, and fluorescence emission was collected between 517 and 571 nm. The mean intracellular fluorescence intensity of one to two cells per root was measured using FIJI (*71*).

### N⁺-BODIPY staining and lifetime imaging

Five-day-old seedlings were stained for 15 min with 10 µM N⁺-BODIPY, washed twice in liquid ½ MS medium, and mounted in liquid ½ MS medium between a microscope slide and coverslip for imaging. Lifetime imaging of N⁺-BODIPY was performed as previously described (*55*) with minor modifications. Briefly, roots were imaged using a Leica SP8 confocal laser scanning microscope (CLSM) (Leica Microsystems GmbH, Wetzlar, Germany) coupled to FastFLIM PicoQuant system consisting of a Sepia Multichannel Picosecond Diode Laser, a PicoQuant Timeharp 260, a TCSPC Module, and a Picosecond Event Timer. Imaging was performed using a 40x water immersion objective (NA 1.2). N⁺-BODIPY was excited at 440 nm using a pulsed laser (LDH-P-C-470, PicoQuant) with a rate of 40 Hz. Fluorescence emission was collected using an HyD4 SMD from 455 nm to 505 nm with a time-correlated single-photon counting (TCSPC) resolution of 25 ps. FLIM acquisition and integration were performed using PicoQuant SymPhoTime software. One to two plasma membrane regions of interest per seedling were selected for lifetime fitting. Lifetime fitting was performed using a two-component exponential reconvolution model, from which τ₁ was extracted for comparison between conditions.

### Di-4-ANEPPDHQ staining, ratiometric imaging and lifetime imaging

Five-day-old seedlings were stained for 5 min with 5 µM di-4-ANEPPDHQ (ThermoFisher Scientific, D36802), washed three times in liquid ½ MS medium, and mounted in liquid ½ MS medium between a microscope slide and coverslip for imaging. For ratiometric measurements, seedlings were imaged using an inverted Zeiss LSM 880 confocal microscope equipped with a 40x water-immersion objective (NA 1.2). Di-4-ANEPPDHQ was excited at 488 nm, and fluorescence emission was collected between 560-590 nm and 640-670 nm. Ratiometric images were generated and analyzed in FIJI (*71*). For fluorescence lifetime imaging, seedlings were imaged using an inverted Olympus FV3000 microscope equipped with a 60x water-immersion objective (NA 1.2) and a filter cube for GFP/RFP. Excitation was performed using a pulsed laser (40 Hz repetition rate) at 485 nm, and fluorescence was collected with a photon-counting PMA hybrid 40 detector using a scanning resolution of 512 × 512 pixels and a TCSPC resolution of 25.0 ps until 1,000 photons per pixel were collected. FLIM acquisition and integration were performed using PicoQuant SymPhoTime software. One to two plasma membrane regions of interest per seedling were selected for lifetime fitting. Lifetime fitting was performed using a two-component exponential reconvolution model, from which τ₁ was extracted for comparison between conditions.

### Analysis of intracellular ROS accumulation

Five-day-old seedlings expressing the HyPer7 sensor (*57*) were imaged using a Leica SP8 confocal microscope equipped with a 40x water-immersion objective (NA 1.2). Two excitation wavelengths were used, 488 nm and 405 nm, corresponding to the oxidized and reduced forms of the probe, respectively. Fluorescence emission was collected between 508 and 535 nm. Ratiometric images were generated using the “Divide” option of the Image Calculator function in FIJI (*71*) and mean fluorescence intensity was measured in one to two regions of interest per seedling.

### Analysis of intracellular Ca^2+^ and Cl^-^

Intracellular Ca^2+^ and Cl^-^ accumulation were analyzed using the 2-in-1 genetically encoded fluorescent indicator R-GECO1-GSL-E^2^GFP (*42*). For observations made after 30 min of treatment, five-day-old seedlings were mounted in liquid ½ MS medium containing 10 µM DSF or the corresponding mock solution (0.0001% DMSO). For observations made after 24 hours of treatment, five-day-old seedlings were mounted on a block of solid ½ MS medium containing 10 µM DSF or the corresponding mock solution (0.0001% DMSO) and placed in a homemade chamber for imaging. Seedlings were imaged using a Leica SP8 confocal microscope equipped with a 40x water-immersion objective (NA 1.2). R-GECO1 was imaged using excitation at 561 nm and fluorescence emission was collected between 580–630 nm. E_2_GFP was excited at 458 nm and fluorescence emission was collected between 500–550 nm which corresponds to its pH isosbestic point (*42*, *72*).

### Aequorin-based analysis of intracellular Ca^2+^

The analysis of rapid calcium influx upon elicitation was performed as previously described (*27*). Briefly, leaf discs (4 mm diameter) were sampled from 5-week-old Col-0/Aequorin (*73*) plants, placed into the wells of a 96-well plate containing 100 µL of 125 µM coelenterazine H (MedChemExpress, HY-D1024) and incubated overnight in the dark. The following day, coelenterazine H was removed in the dark and replaced with 100 µL of double-distilled water. Luminescence was recorded for 10 min, after which 25 µL of microbial-derived molecules at 5x concentration was added to each well, and luminescence was recorded for an additional 45 min using a GloMax plate reader (Promega). Cytosolic Ca^2+^ concentration is expressed as *L*/*L_ma_*_x_, with *L* corresponding to the photons collected at each time point and *L_max_*corresponding to the photons collected during a 10 min discharge of the remaining aequorin using 100 µL of 2 M CaCl₂ in 20 % ethanol. For long-term measurement of DSF-induced calcium accumulation, leaf discs (4 mm diameter) were sampled from 5-week-old Col-0/Aequorin plants and placed into the wells of a 96-well plate containing 100 µL of 125 µM coelenterazine H supplemented with 10 µM DSF or the corresponding control solution (0.0001% DMSO) and incubated in the dark overnight. The following day, the coelenterazine H solution was removed in the dark and replaced with 300 µL water. Luminescence was then recorded for 4 hours, starting from 16 to 20 hours after DSF treatment, using a GloMax plate reader (Promega).

### RNA isolation and sequencing

Leaves of four-week-old plants were infiltrated with 40 µM DSF or an equivalent dilution of DMSO, sampled after 30 min or 24 hours of treatment in a 2 mL tube containing a ceramic bead and were immediately flash frozen in liquid nitrogen. RNA isolation was conducted as described by Albrecht et al. (2012). Briefly, frozen samples were ground using a TissueLyser (BioRad) and vortexed with TRIzol™ Reagent (ThermoFisher scientific) and chloroform to extract RNA. RNA was precipitated using isopropanol and the recovered RNA pellet was washed with ethanol and resuspended in sterile double-distilled water. RNA samples were then cleaned using the RNA Clean & Concentrator-25 kit (Zymo, R1017) following manufacturer’s recommendations. Sample quality control and RNA sequencing were conducted by BMKGENE (Biomarker Technologies GmbH). The samples were sequenced on the Illumina NovaSeq X platform. Raw reads were uploaded and processed on Galaxy (https://usegalaxy.eu/). Paired reads were trimmed using the Trimmomatic tool with a sliding window of 4 bp and a quality threshold set to 20. Quality control was performed using FastQC and estimated counts were obtained using the Kallisto pseudo-alignment method against the reference transcriptome Col-0 TAIR10.

### Transcriptomic analysis

Transcriptomic analysis was conducted from gene-level estimated counts using the DESeq2 package (Love et al., 2014) in RStudio 2023.09.1+494 (http://www.rstudio.com/). Count normalization, dispersion estimation, and model fitting were performed according to the standard DESeq2 pipeline. Statistical significance was evaluated using the Wald test implemented in DESeq2. Genes with a p-value < 0.05 and an absolute log2 fold change greater than 1 were defined as significantly differentially expressed genes. For the comparison of DSF-responsive genes with published elicitor-responsive transcriptomes from flg22, elf18, Pep1, nlp20, OGs, ch8, and LPS (*43*), genes were scored as DSF- or elicitor-induced when the log2 fold change was ≥ 1, considering either individual time points or all time points combined, and overlaps were visualized using UpSet plots (ComplexUpset and UpSetR packages). For transcriptomic comparisons of DSF-, PTI-, and ETI-responsive genes, PTI was represented by the flg22 dataset, while the ETI-responsive gene list was obtained using an effector-induced published dataset (Ngou et al., 2021) filtered for *P* value ≤ 0.05 and absolute log2 fold change ≥ 1. Integrated DSF, flg22, and ETI expression values were displayed as row-scaled heatmaps (pheatmap package) and intersections between DSF, PTI and ETI gene sets were visualized using Venn/Euler diagrams (ggVennDiagram package). The list of commonly considered immune responsive genes was curated from Arabidopsis immunity literature covering cell-surface receptors, intracellular NLRs, and downstream kinase signaling modules (*74–76*), whereas the list of hormonal response genes was assembled from reviews and primary studies on defense hormone pathways and systemic immune small molecules (*77–80*).

### Apoplastic ROS measurement

The analysis of apoplastic ROS production was performed as previously described (*81*). Briefly, leaf discs (4 mm diameter) were collected in 96-well plates containing sterile water and incubated overnight. The next day the water was replaced with a solution containing 0.5 mM L-012 (FUJIFILM Wako Pure Chemical Corporation, 120-04891), 20 μg/mL horseradish peroxidase (HRP, Merck, P8375), DSF, flg22, 3-OH-FA or their corresponding mock controls at the indicated concentration. Luminescence was recorded for at least 45 min using a GloMax plate reader (Promega). To test the effect of DSF on flg22-induced ROS, leaf discs were pre-treated with DSF at 5 or 40 µM or the corresponding control solvent (DMSO) for the indicated time before sample collection.

### Pto DC3000 infection assay

Five-week-old plants were infiltrated with 5 µM or 40 µM DSF or 1 µM flg22 or the corresponding control solution for 24 hours. *Pseudomonas syringae* pv. *tomato* (Pto) DC3000 cells were streaked out from glycerol stock on fresh King’s B medium plates containing 1 % agar supplemented with 50 μg/mL rifampicin. Bacteria were collected from plates and resuspended in liquid King’s B medium to grow overnight at 28°C. The overnight culture was used to inoculate fresh King’s B medium and grown for 1 hour. After centrifugation of the cells and resuspension in MgCl₂, the bacterial solution was adjusted to 10⁵ colony forming unit (cfu)/mL of *Pst* DC3000 to be infiltrated in leaves. Three days post-inoculation, two leaf discs (4 mm diameter) were sampled per leaf and ground in 200 µL of sterile 10 mM MgCl₂. Serial dilutions were prepared from the homogenate, plated on King’s B medium and colonies were counted two days later.

### Protein extraction and western blotting

Five-day-old seedlings grown on solid ½ MS medium were transferred to liquid ½ MS medium, where they were grown for an additional 7 days. Following the treatments specified in the corresponding figures, seedlings were flash-frozen in liquid nitrogen and ground in protein extraction buffer containing 150 mM NaCl, 10% glycerol, 50 mM Tris-HCl (pH 7.5), 1% IGEPAL, 100 µM phenylmethylsulfonyl fluoride, 0.2 mM 2-aminoethyl benzenesulfonyl fluoride, 0.7 µM bestatin hydrochloride, 0.7 µM pepstatin A, 10 µM leupeptin hydrochloride, 1.4 µM E-64, 1.4 µM phenanthroline, 5 mM DTT, 2.5 mM NaF, 1 mM Na₂MoO₄, and 2 mM Na₃VO₄. Samples were incubated for at least 45 min and then centrifuged at 14,000 x g for 15 min. The supernatant was collected and boiled in Laemmli buffer at 95 °C for 5 min. Proteins were separated using a 10 % SDS-PAGE and transferred to a PVDF membrane activated in methanol. After blocking in milk dissolved in TBS 1% with 0.1% Tween 20, the blots were probed with 1:3000 anti-pERK42-44 (Cell Signaling; 9381) diluted in 5 % BSA for phosphorylated pMAPK or with 1:5000 anti-BAK1 (Agrisera, AS12 1858) or 1:5000 anti-FLS2 (Agrisera, AS12 1857) in 5 % milk. After washing and treatment with SuperSignal West Pico PLUS (ThermoFisher), the blots were imaged using the ImageQuant LAS 4000 mini (GE Healthcare Life Sciences).

### Statistical analysis

The number of independent experiments, the number of individual plants, as well as the number of cells analyzed when applicable are reported in each figure legend. Data analyses and statistical tests were performed using R 4.3.1 (https://cran.r-project.org/) and RStudio 2023.09.1+494 (http://www.rstudio.com/). The statistical tests used are specified in the figure legends.

---

**Figure S1.**
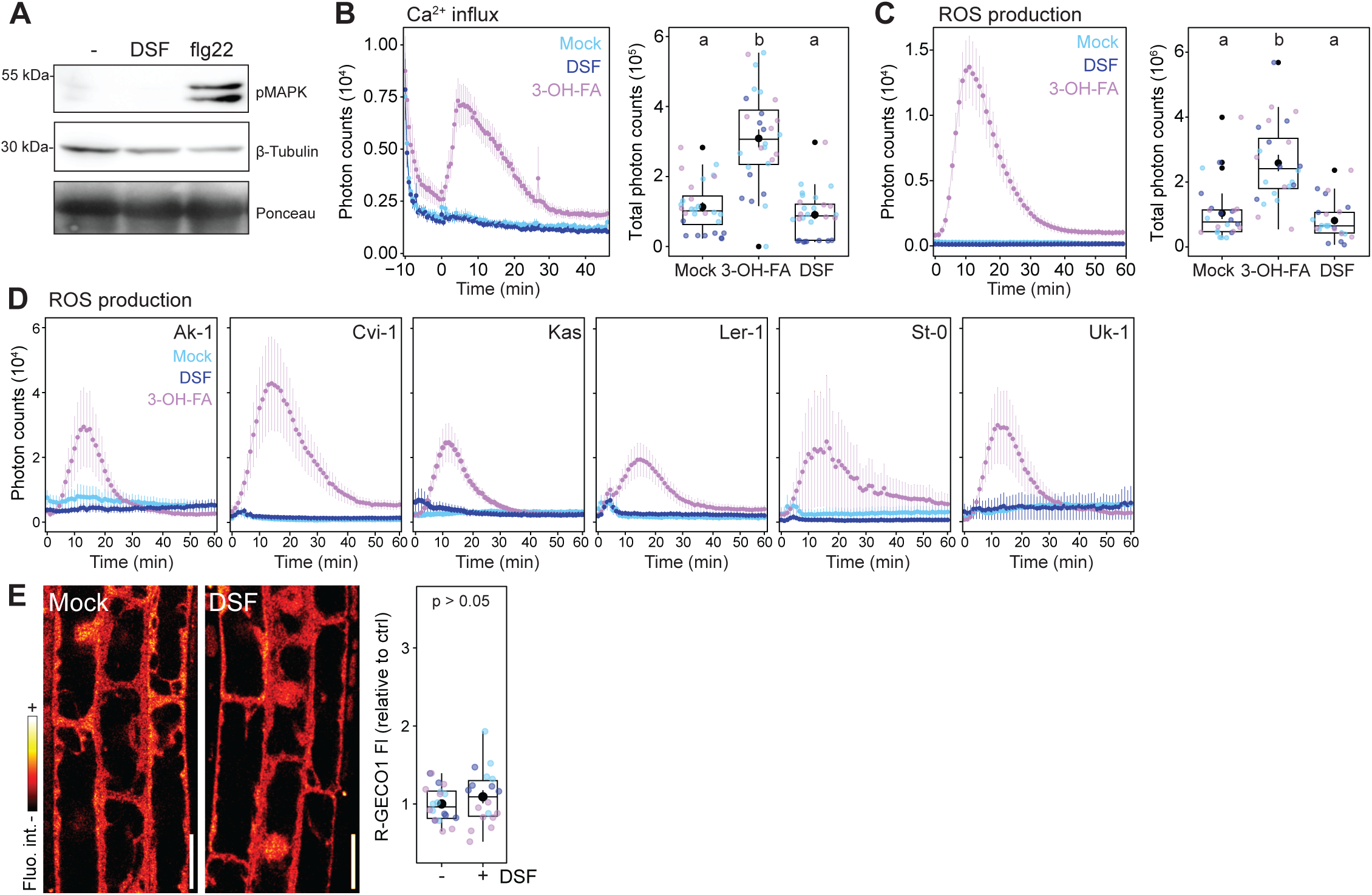
DSF does not induce hallmarks of PTI signaling in Arabidopsis. (**A**) Western blot analysis of MAPK phosphorylation in Arabidopsis seedlings treated for 10 min with 10 µM DSF or 100 nM flg22. Membranes were probed with anti-pERK42/44 or β-Tubulin antibodies to assess phosphorylated MAPK or protein loading. (**B**) Time course analysis of Ca^2+^ influx in Col-0/Aequorin leaf discs treated with 5 µM 3-OH-FA or 40 µM DSF. The boxplots represent total photon counts (0-45 min). Each data point represents a leaf disc; colors represent independent experiments with 24 leaf discs analyzed per condition. Different letters indicate statistically significant differences between conditions (one-way ANOVA followed by a Tukey’s HSD; p<0.01). (**C**) Time course analysis of apoplastic ROS production after treatment with 5 µM 3-OH-FA or 40 µM DSF or corresponding mock (DMSO) in Col-0. Bars represent the standard error. The boxplots represent the mean total photon counts (0-60 min). Each data point represents a leaf disc; colors indicate independent experiments with 24 leaf discs analyzed per condition. Different letters indicate statistically significant differences between conditions (one-way ANOVA followed by a Tukey’s HSD; p<0.01). (**D**) Time course analysis of apoplastic ROS upon treatment with 5 µM 3-OH-FA or 40 µM DSF or corresponding mock control (DMSO) in Arabidopsis ecotypes. Bars represent the standard error. (**E-F**). Analysis of intracellular Ca^2+^. Representative confocal images (E) and mean fluorescence intensity (F) of R-GECO1 in Col-0/R-GECO1-GSL-E^2^GFP root epidermal cell treated with 10 µM DSF or mock (DMSO) for 30 min. Each data point represents the mean intracellular fluorescence per cell; colors represent independent experiments with at least 10 seedlings analyzed per condition. *P* value reports a Wilcoxon test. Scale bar = 20 µm.

**Figure S2.**
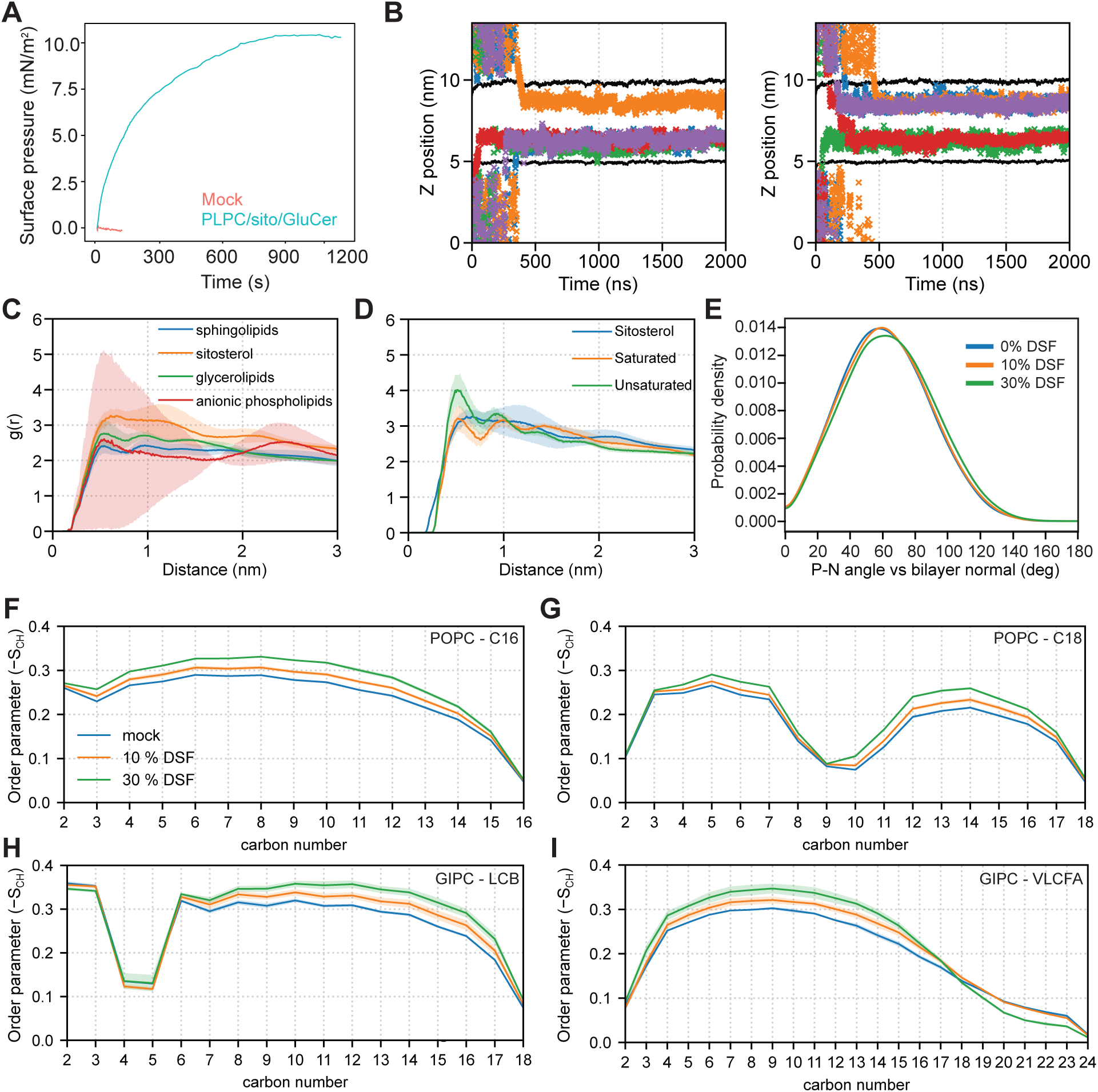
Analyses of DSF insertion in PM model membranes. (**A**) Adsorption kinetics of DSF (12 μM in the subphase 10 mM MES, 150 mM NaCl, pH 5.8) into PLPC/sitosterol/GluCer (6:2:2 molar ratio) monolayer at an initial surface of 15.0 ± 0.4 mN/m. Z position of DSF center of mass over the time for simulation repeats not shown in Fig 1D. Each cross represents an individual DSF molecule at each time point. Different colors represent five different DSF molecules. Average z position of the phosphorus headgroup atoms is shown as a black line. (**C**) Radial distribution functions *g(r)* ± sd of major lipid classes around DSF. (**D**) Radial distribution functions *g(r)* ± sd of sitosterol and acyl-chain saturation classes around DSF. (**E**) Distribution of the angle between the phospholipid P-N vector and the bilayer normal in the absence and presence of DSF (10%, 30%). (**F**-**I**) Lipid chain order parameters in the absence (in blue) and presence of DSF 10% (in orange), and 30% (in green). Carbon-specific (−SCH) profiles are shown for (F) POPC-C16, (G) POPC-C18, (H) GIPC-LCB, and (I) GIPC-VLCFA.

**Figure S3.**
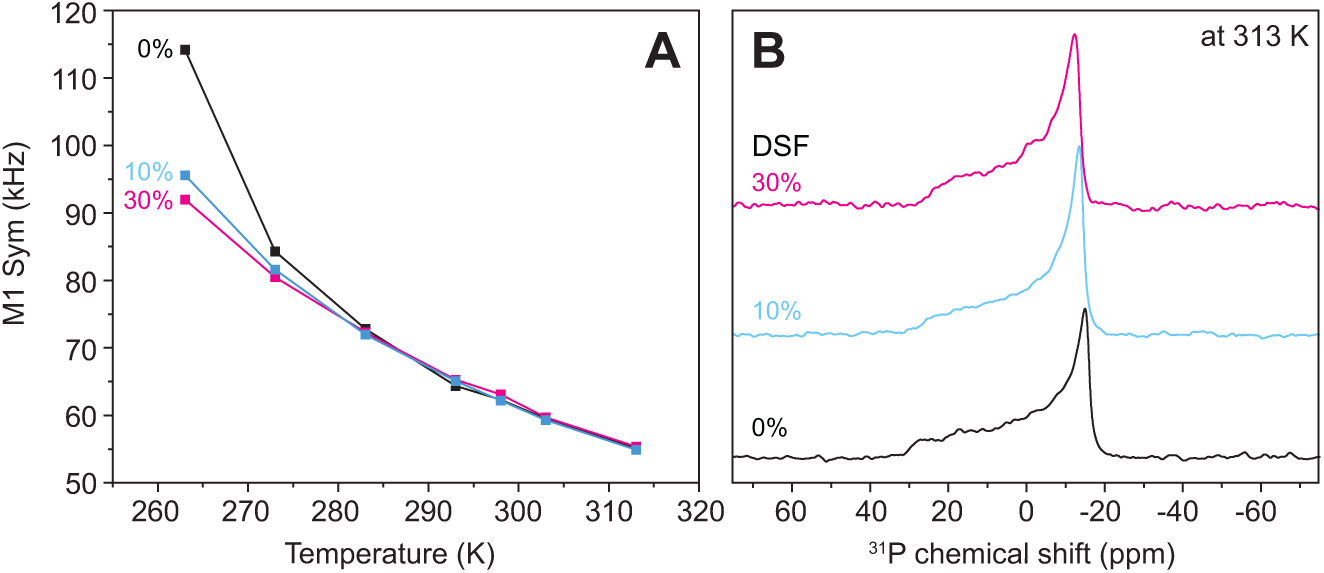
Solid-state NMR analyses at different temperatures. Solid-state NMR spectroscopy of POPC-d31/sitoterol/Cer 7:1:2 (molar ratio) liposomes in the absence (black) or in presence of 10 % (light blue) or 30 % (dark blue) DSF, (**A**) first spectral Moment M_1_ derived from ^2^H quadrupolar echo spectra (**B**) ^31^P Hahn echo spectra detecting the linewidth of the chemical shift anisotropy at 313K.

**Figure S4.**
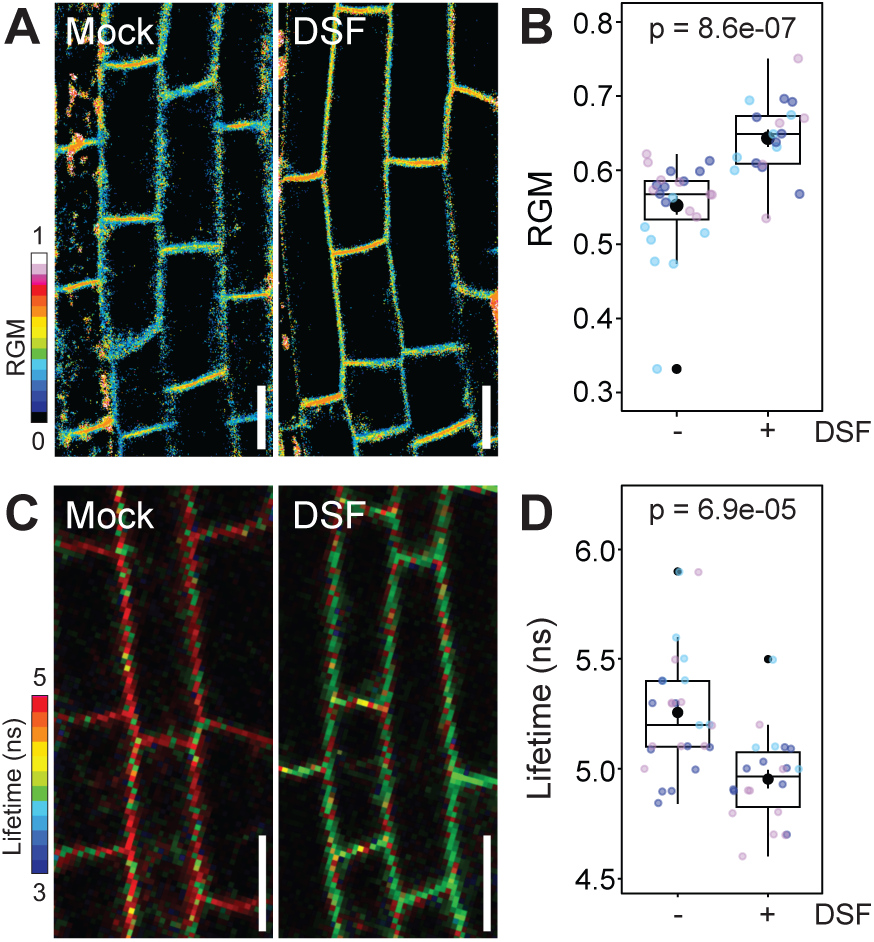
Analysis of plasma membrane order upon DSF treatment. (**A**-**B**) Analysis of plasma membrane order using the di-4-ANEPPDHQ dye. Representative confocal images of di-4-ANEPPDHQ (A) and ratiometric analysis of the mean fluorescence intensity (B) collected between 640-670 nm (red) and 560-590 nm (green), termed RGM which reports on plasma membrane order, in root epidermal cells treated with 40 µM DSF or mock (DMSO) for 30 min. Each data point represents a seedling; colors represent independent experiments with at least 19 seedlings analyzed per condition. *P* value reports a Wilcoxon test. Scale bar = 20 µm. (**C**-**D**) Analysis of plasma membrane microviscosity using the N^+^-BODIPY dye. Representative FLIM images of N^+^-BODIPY (C) and analysis of the mean lifetime (τ1) (D) in root epidermal cells treated with 40 µM DSF or mock (DMSO) for 30 min. Each data point represents a seedling; colors represent independent experiments with at least 14 seedlings analyzed per condition. *P* value reports a Wilcoxon test. Scale bar = 20 µm.

**Figure S5.**
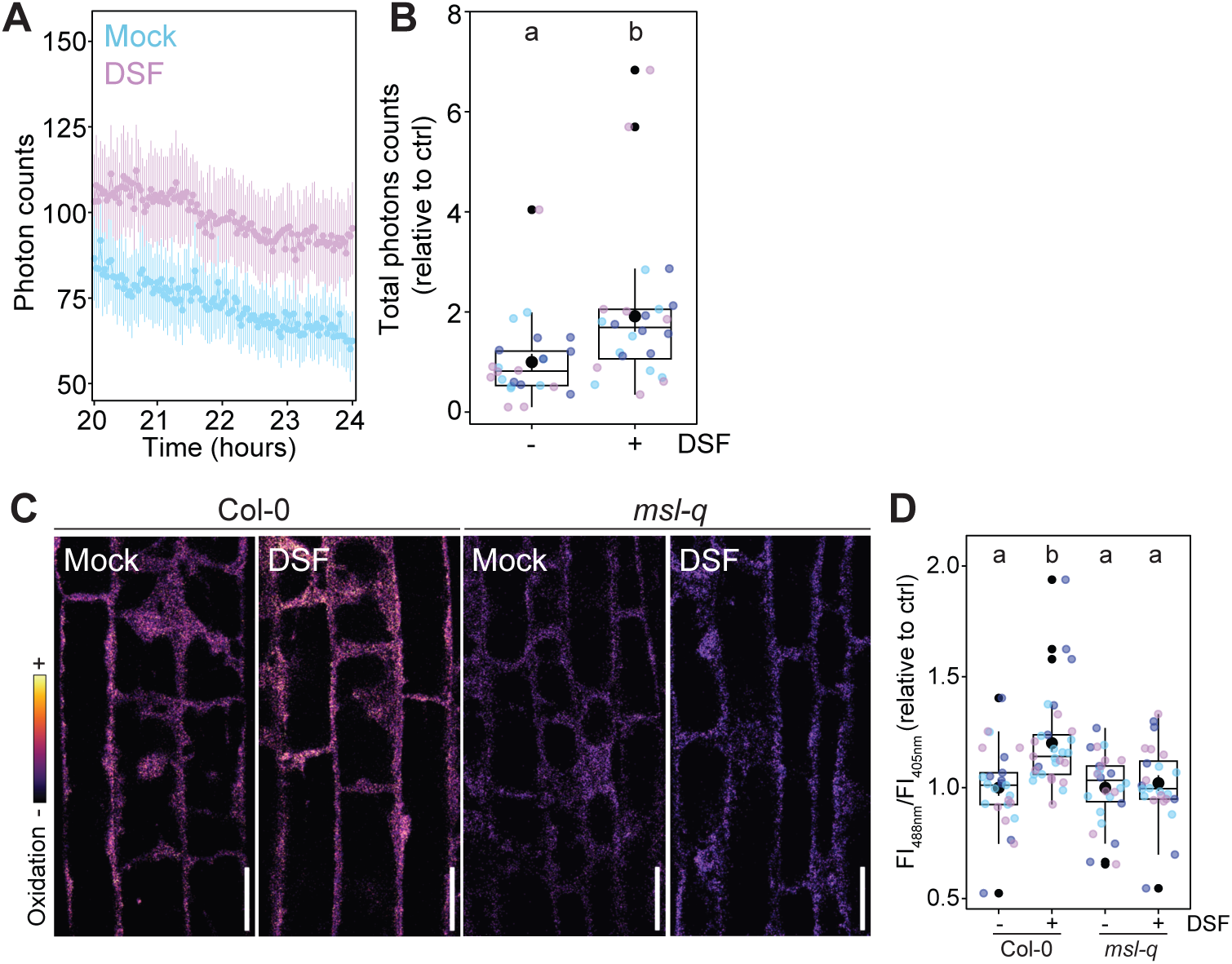
Prolonged DSF treatment induces the accumulation of cytoplasmic Ca^2+^ and a sustained accumulation of intracellular ROS. (**A**-**B**). Aequorin-based analysis of cytoplasmic Ca^2+^. Time course (A) and total photon counts (B) analysis of cytoplasmic Ca^2^ in Col-0/Aequorin leaf discs treated with 10 µM DSF or mock (DMSO). The boxplot represents the mean total photon counts from a 4 h interval. Each data point represents a leaf disc; colors represent independent experiments with 24 leaf discs analyzed per condition. Different letters indicate statistically significant differences between conditions (p<0.0001; Mann-Whitney test). (**C**-**D**) Analysis of intracellular ROS. Representative confocal images (C) and ratiometric analysis of fluorescence intensity (D) of Hyper7 in root epidermal cells treated with 10 µM DSF or mock (DMSO) for 24 h. Each data point represents mean fluorescence of per seedling; colors indicate independent experiments with at least 22 seedlings analyzed per condition. Different letters indicate statistically significant differences between conditions (one-way ANOVA followed by a Tukey’s HSD; p<0.001).

**Figure S6.**
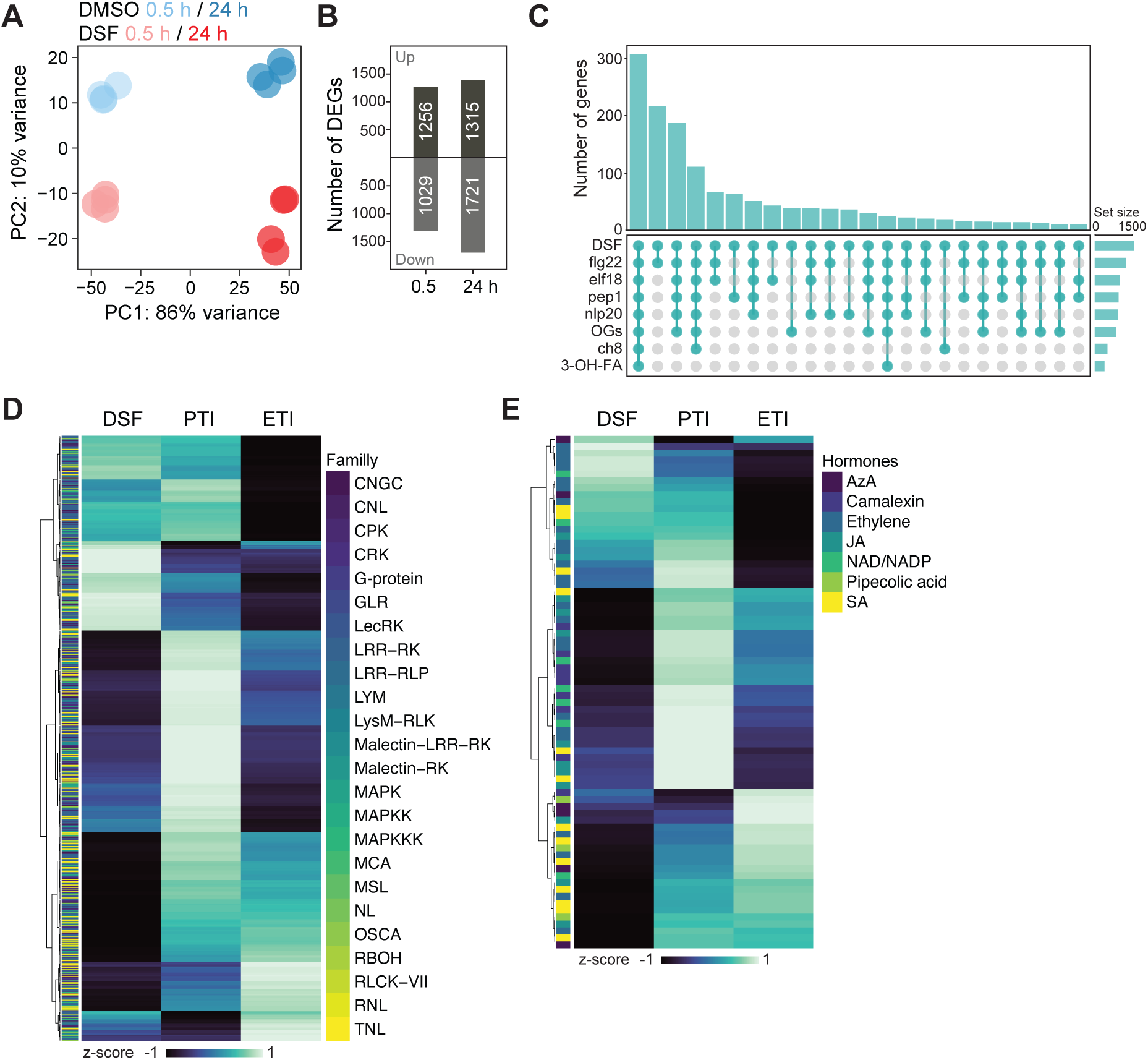
Analysis of DSF-induced changes in transcriptome. (**A**) Principal component analysis of transcriptomes from mock-treated (DMSO) and DSF-treated Col-0 plants for 0.5 h and 24 h. (**B**) Number of significantly upregulated and downregulated genes Col-0 plants treated for 0.5 h and 24 h. (**C**) Overlap of DSF-responsive genes with elicitor-induced genes, shown as an UpSet plot. (**D-E**) Heatmap of DEGs commonly associated with immune responses (D) or with hormonal pathways (E) induced by DSF, MAMPs (PTI), and AvrRps4 (ETI), normalized as z-scores.

**Figure S7.**
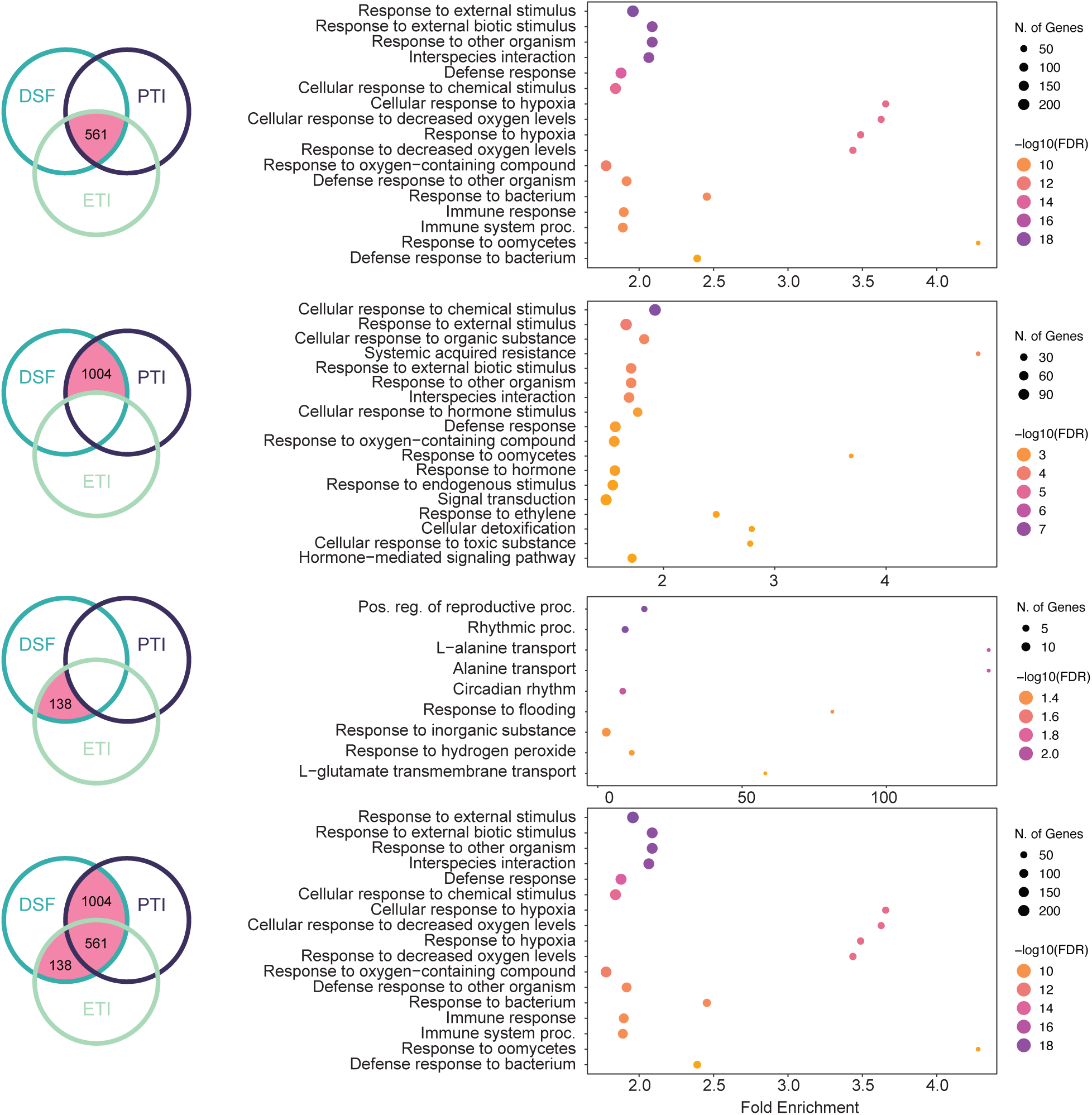
Gene ontology analyses of DEGs common to DSF, ETI and PTI. The intersections (left) tested for Gene Ontology enrichment (right) are marked in pink.

**Figure S8.**
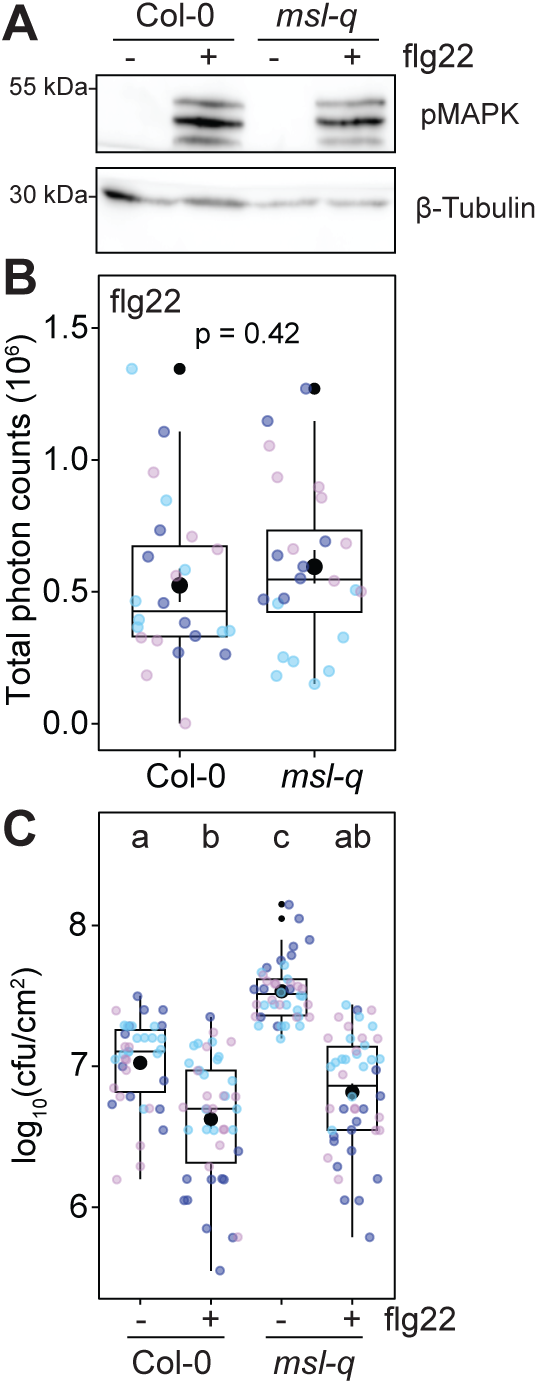
***MSLs* are not required for flg22-induced signaling and disease resistance.** (**A**) Western blot analysis of MAPK phosphorylation in Arabidopsis seedlings treated for 10 min with 100 nM flg22. Membranes were probed with anti-pERK42/44 or β-Tubulin antibodies to assess phosphorylated MAPK or protein loading. (**B**) Apoplastic ROS production after 100 nM flg22 in Col-0 and *msl-q*. Each data point represents an individual leaf disc; colors indicate independent experiments with 24 leaf discs analyzed per condition. *P* value reports a Wilcoxon test. (**C**) Number of *Pto* DC3000 bacteria 3 days after inoculation of plants pre-treated with 5 µM DSF or mock (DMSO) for 24 h. Each data point represents an individual leaf; colors indicate independent experiments. Different letters indicate statistically significant differences between conditions (one-way ANOVA followed by a Tukey’s HSD; p<0.05).

**Figure S9.**
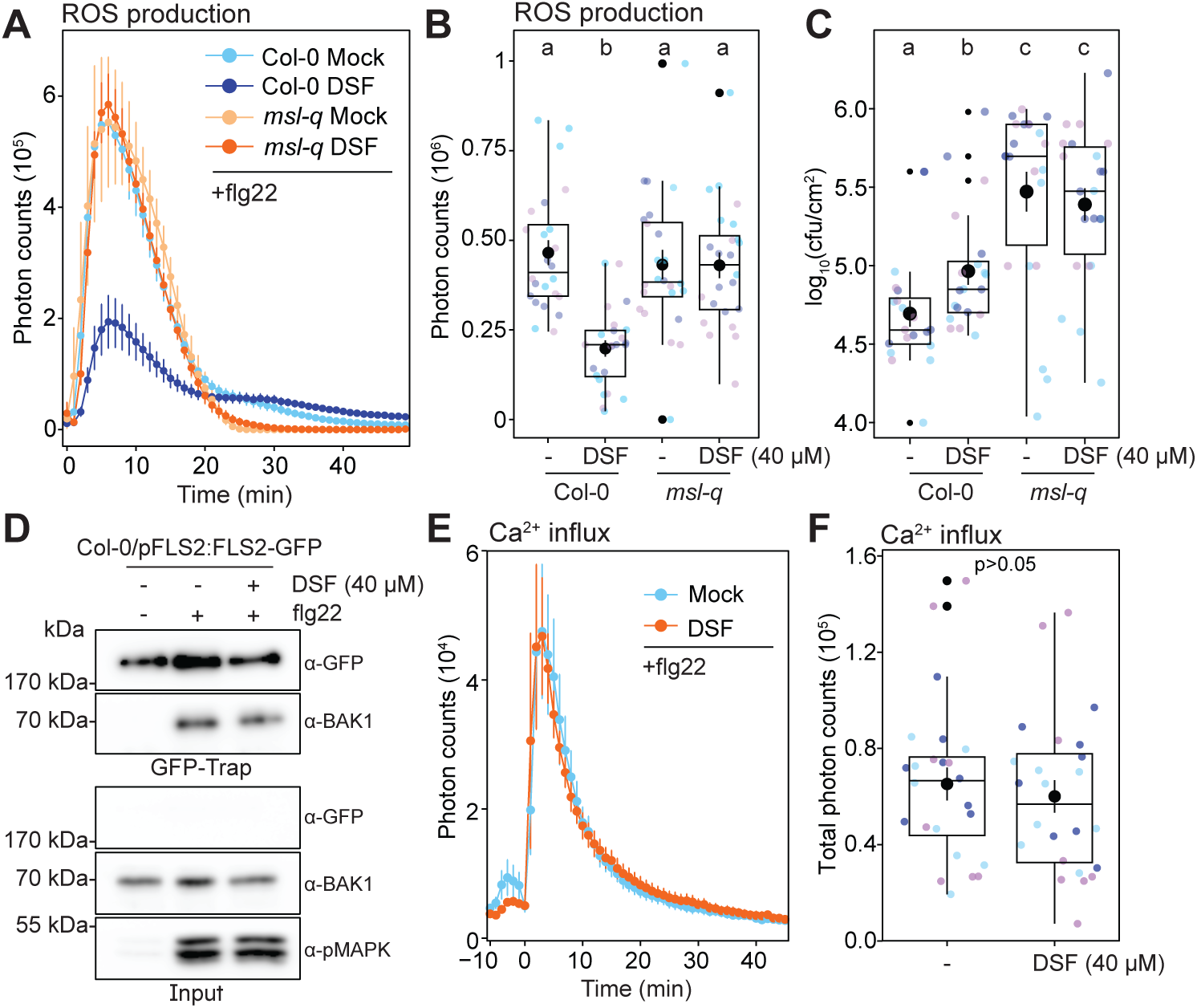
Analysis of DSF 40 µM treatment. (**A-B**) Luminescence-based analysis of apoplastic ROS production upon elicitation with 100 nM flg22. Time course (A) and total photon counts (B) analysis of apoplastic ROS in Col-0 and *msl-q* leaf discs treated with 40 µM DSF or mock (DMSO). The boxplot represents the mean total photon counts. Each data point represents a leaf disc; colors represent independent experiments with 24 leaf discs analyzed per condition. Different letters indicate statistically significant differences between conditions (one-way ANOVA followed by a Tukey’s HSD). (**C**) Number of *Pto* DC3000 bacteria 3 days after inoculation of plants pre-treated with 40 µM DSF or mock (DMSO) for 24 h. Each data point represents an individual leaf; colors indicate independent experiments. Different letters indicate statistically significant differences between conditions (one-way ANOVA followed by a Tukey’s HSD; p<0.05). (**D**) Co-immunoprecipitation in Col-0/pFLS2:FLS2-GFP pre-treated with 40 µM DSF or mock (DMSO) for 24 h and elicited with 100 nM flg22 for 10 min. Western blots were probed with α-FLS2, α-BAK1 or α-pER42 44 antibodies to detect phosphorylated MAPK. (**E-F**) Aequorin-based analysis of cytoplasmic Ca^2+^. Time course (E) and total photon counts (F) analysis of cytoplasmic Ca^2^ in Col-0/Aequorin leaf discs pre-treated with 40 µM DSF or mock (DMSO) for 24 h and elicited with 100 nM flg22. The boxplot represents the mean total photon counts (0-45 min). Each data point represents a leaf disc; colors represent independent experiments with 24 leaf discs analyzed per condition. *P* value reports a Wilcoxon test.

## Notes

### Competing Interest Statement

The authors have declared no competing interest.

### Summary of Updates

The manuscript has been further proofread

